# Accumulating computational resource usage of genomic data analysis workflow to optimize cloud computing instance selection

**DOI:** 10.1101/456756

**Authors:** Tazro Ohta, Tomoya Tanjo, Osamu Ogasawara

## Abstract

**Background:** Container virtualization technologies such as Docker became popular in the bioinformatics domain as they improve portability and reproducibility of software deployment. Along with software packaged in containers, the workflow description standards Common Workflow Language also enabled to perform data analysis on multiple different computing environments with ease. These technologies accelerate the use of on-demand cloud computing platform which can scale out according to the amount of data. However, to optimize the time and the budget on a use of cloud, users need to select a suitable instance type corresponding to the resource requirements of their workflows.

**Results:** We developed CWL-metrics, a system to collect runtime metrics of Docker containers and workflow metadata to analyze resource requirement of workflows. We demonstrated the analysis by using seven transcriptome quantification workflows on six instance types. The result showed instance type options of lower financial cost and faster execution time with required amount of computational resources.

**Conclusions:** The summary of resource requirements of workflow executions provided by CWL-metrics can help users to optimize the selection of cloud computing instance. The runtime metrics data also accelerate to share workflows among different workflow management frameworks.

## Background

According to the improvement of DNA sequencing technology in accuracy and quantity, various sequencing methods are now available to measure different genomic features. Each method produces a massive amount of nucleotide sequence data that requires a different data processing approach [1]. Bioinformatics researchers develop data analysis tools for each sequencing technique, and they publish implementations as open source software [2]. To start data analysis, researchers need to select the tools by their experimental design and install them to their computing environment.

Installing open source tools in one’s computational environment is, however, not always straightforward. Tools developed by different developers and different programming framework require different prerequisites, which forces one to follow the instruction provided by each tool’s developer. Installing various software in one environment also can occur a conflict of software dependencies that are hard to resolve. Even if one could successfully install all the tools required for the analysis, maintaining the environment where all the tools keep working as expected is also a burden. There are also many events that can break the environment such as changes or updates of hardware, operating system, or software libraries. Therefore, the complexity of data analysis environment management gets higher when a project performs genomic data analysis that requires many tools. The high cost of setting up an environment results in the prevention of scaling out the computational resources as well. The difficulty also brings researchers’ dependency to the existing computing platform already set up, and the concentration of data processing jobs to the limited resource.

The container virtualization technology, represented by Docker, enables users to create a software runtime environment isolated from the host machine [3]. This technology that is getting popular also in the biomedical research domain is a promising method to solve the problem of installing software tools [4]. Along with the containers, using workflow description and execution frameworks such as those from the Galaxy project [5] or the Common Workflow Language (CWL) project [6] lowered the barrier to deploy the data analysis environment to a new computing environment. Moreover, the workflows described in a standardized format can help researchers to share the environment with collaborators with ease. The improvement of portability of data analysis environment, consequently, has made the on-demand cloud infrastructure an appealing option for researchers.

On-demand cloud is beneficial for most cases in genome science because users can increase or decrease the number of computing instances without maintaining hardware as the amount of data from laboratory experiments changes [7]. For example, some sequencing applications require data analysis software that uses a considerable amount of memory, but individual research projects often cannot afford such a large scale computing platform. Users can save their budget by using the on-demand cloud platform as most of the service providers charge per usage.

However, to use an on-demand cloud environment efficiently regarding time and economic cost, it is essential to select a suitable computing unit, so-called instance type, from many options offered by the cloud service providers. For example, Amazon Web Service (AWS), one of the popular cloud service providers, offers instance types of different scales for five categories (general purpose, compute optimized, memory optimized, accelerated computing, and storage optimized) [8]. Each data analysis tool has the different minimum requirement of computational resources such as memory or storage, and it can change by input parameters. Executing data analysis workflows on an instance without enough computational resource will result in a runtime failure or unexpected outputs. For example, tools to assemble short reads to construct genome by constructing De Bruijn graph usually take long processing time and a large amount of memory. If one failed to estimate the required amount of memory, the process might fail after a few days of execution, which results in losing one’s time and budget. Thus, users need to know the minimum amount of computational resource required by the execution of their workflows to select a suitable instance type.

To optimize the instance type selection concerning processing time or running cost, users need to compare runtime metrics of workflow executions on environments of different computational specs. Here, we developed CWL-metrics, a system to accumulate runtime metrics of workflow executions with information of the workflow and the machine environment. CWL-metrics provides runtime metrics summary such as usage of CPU, memory, storage I/O with workflow’s input files and parameters to help users to select the proper cloud instance for their workflows.

## Results

### Implementation of CWL-metrics

CWL-metrics is designed to capture runtime metrics data of workflows described in CWL, a workflow description specification developed by an open source community. We designed the system as it does not require the users to perform any configurations to capture runtime metrics. Figure 1 shows the procedures of runtime metrics collection by CWL-metrics. To start collecting metrics, one only needs to install the system, and then run their workflows with cwltool, a reference implementation of CWL [9]. After the installation, the system starts monitoring the processes running on the host machine. Once the system found a cwltool process, it automatically starts collecting runtime metrics via Docker API and environmental information from the host machine. CWL-metrics also captures the log file generated by cwltool to extract workflow metadata such as input files and input parameters.

**Figure 1:**
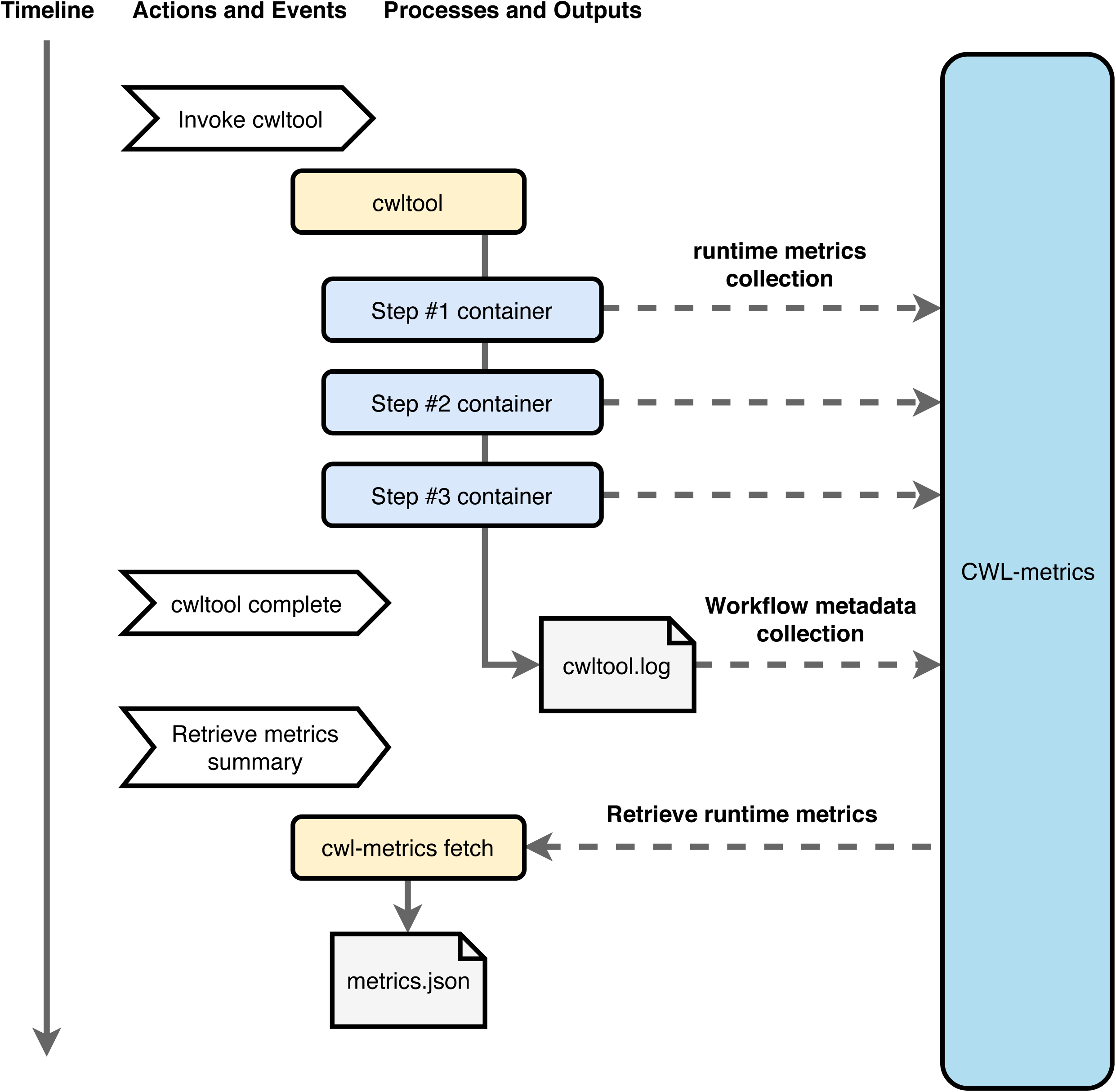
The container runtime metrics collection procedure with CWL-metrics. CWL-metrics was designed to capture runtime metrics of workflow steps automatically. After the initialization of the system, users only need to run a workflow by cwltool to start metrics capturing. The system collects runtime metrics of containers, and then the workflow metadata is captured after the workflow process finished. To retrieve runtime metrics, using the *cwl-metrics* command can output summary data in JSON or tab-delimited format.

To capture and store the information from multiple data source, CWL-metrics launches multiple components as Docker containers (Figure 2). These components keep running on the host machine after the initialization to cooperate the data collection. The Telegraf container collects runtime metrics data from the Docker API for every sixty seconds, and send the data to the Elasticsearch container. The Elasticsearch container provides data storage and the data access API. CWL-metrics automatically launches and stops these components on the single host machine. If users need to collect metrics of workflows running on multiple instances, they need to install CWL-metrics on each instance and assemble the summary data after the metrics data capture. Users can use their Elasticsearch server by setting environment variable ES_HOST and ES_PORT before initializing CWL-metrics.

**Figure 2:**
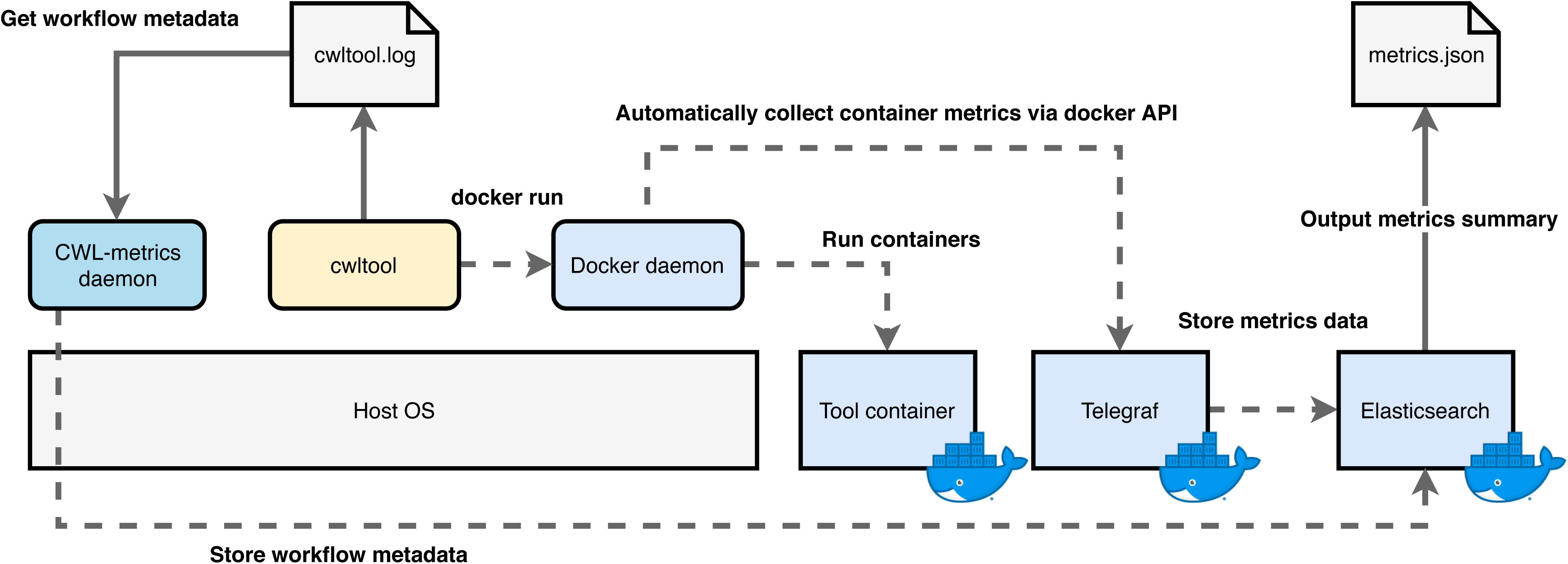
The CWL-metrics components and working process. CWL-metrics runs a daemon process and Docker containers on the host machine. The process and containers keep running until the system is terminated. Once a cwltool process starts running on the same machine, CWL-metrics system monitors the process to get the list of workflow step containers and log files. Every sixty seconds, the Telegraf container try to access the Docker daemon to get runtime metrics of running containers. Fluentd container (not shown in the figure) sends runtime metrics data collected by Telegraf to the Elasticsearch container. CWL-metrics daemon process captures cwltool log file and sends workflow metadata to Elasticsearch.

To access and analyze the data collected by CWL-metrics, users can use the command *cwl-metrics* to get the data in JSON (Figure 3) or tab separated values (TSV) format. The JSON format contains workflow metadata such as the name of the workflow, the time of start and end of the workflow execution. It also has the information of the environment including the total amount of memory and the size of storage available on the machine. The steps field of the JSON format contains information of the runtime metrics, the executed container, and the input files and parameters. Users can parse the data to analyze the performance of a tool execution or the whole workflow. The TSV format provides minimum information for each container execution so that one can easily compare the metrics data of steps.

**Figure 3:**
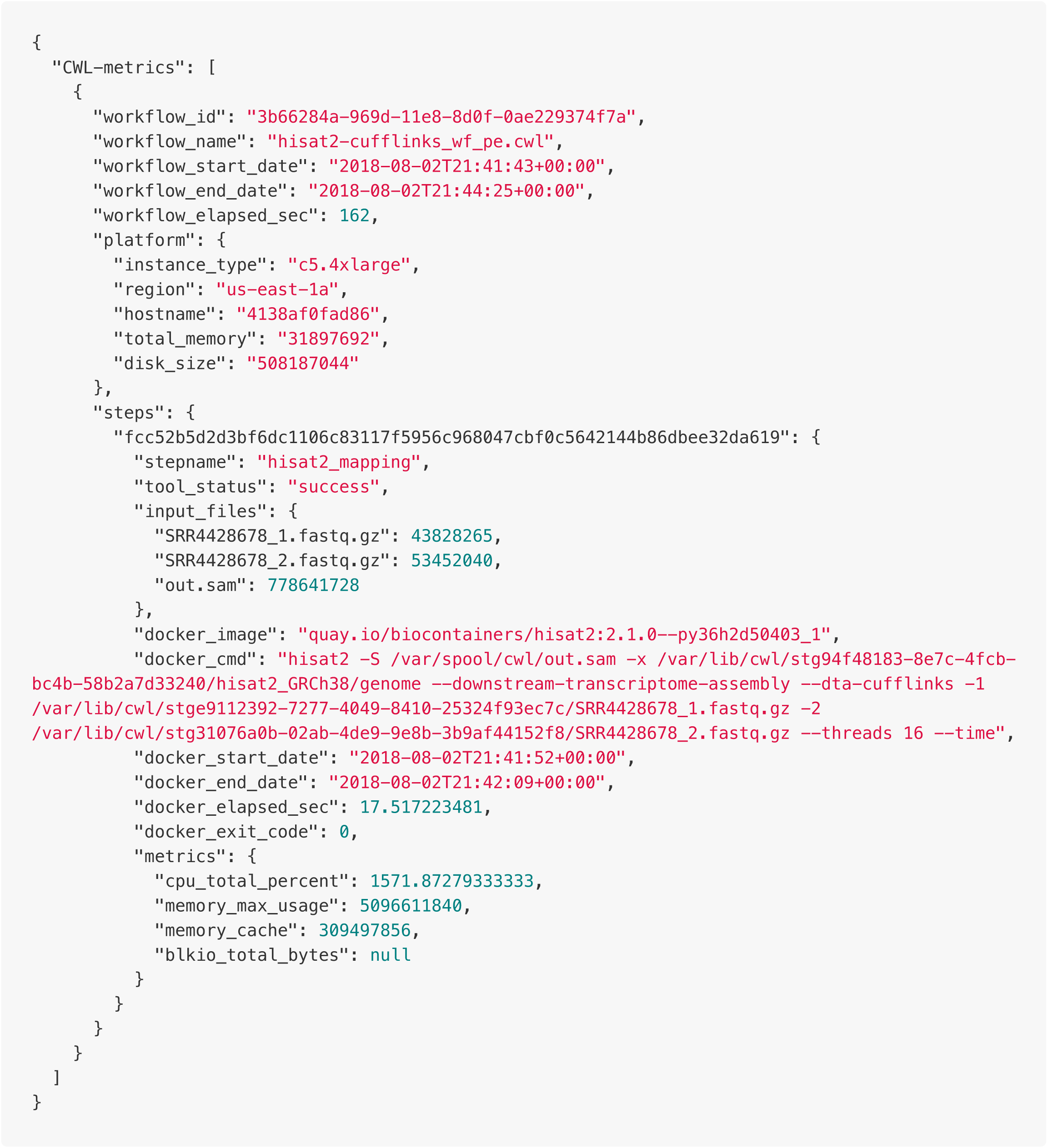
An example of runtime metrics data summarized by CWL-metrics. CWL-metrics can output JSON formatted data which includes workflow metadata, tool container metadata, and tool container runtime metrics. The workflow metadata appears once for one workflow run with data of multiple steps in "steps" key while the example only has one step in the workflow to reduce the number of lines. Each step has a name, exit status, input files with file size, and details of the Docker container. Runtime metric values can be null for short-time steps since CWL-metrics collects these metrics with sixty seconds interval.

### Use CWL-metrics to capture runtime metrics of RNA-Seq workflows

As an example use case to capture and analyze runtime metrics of workflows, we performed an analysis to optimize instance type selection for RNA-Seq quantification workflows. We run seven RNA-Seq workflows (Table 1) for nine public human RNA-Seq data with different read length and number of reads (Table 2) on six types of Amazon Web Service (AWS) Elastic Compute Cloud (EC2) service (Table 3) to capture the runtime metrics by CWL-metrics for each combination. Each workflow description has two different options for read layout; single-end and paired-end. For the selection of workflows, we chose two read mapping tools STAR and Hisat2, with two transcriptome assembly and read count programs Cufflinks and StringTie. We also used two popular tools using alignment-like algorithms, Kallisto and Salmon. TopHat2, the program which was once the most popular, but now obsolete, was added among them for comparing purpose. We performed metrics data collection five times for each combination of workflow, input data, and instance type. The analysis used only the succeeded runs.

**Table 1:** The components of RNA-Seq quantification workflows. We described seven different RNA-Seq quantification workflows in CWL. Each workflow description has two different options for read layout, single-end and paired-end. We selected two major read mapping tools STAR and Hisat2, with two transcriptome assemble and read count programs Cufflinks and StringTie. We also used two popular tools using alignment-like algorithms, Kallisto and Salmon. We added TopHat2, one of the most popular but obsolete program for comparing purpose.

| Workflow name | Steps | CWL definition files |
| --- | --- | --- |
| tophat2-cufflinks | download-sra, pfastq-dump, tophat2-mapping, cufflinks | https://github.com/pitagora-galaxy/cwl/tree/master/workflows/tophat2-cufflinks |
| hisat2-cufflinks | download-sra, pfastq-dump, hisat2-mapping, samtools_sam2bam, samtools_sort, cufflinks | https://github.com/pitagora-galaxy/cwl/tree/master/workflows/hisat2-cufflinks |
| hisat2-stringtie | download-sra, pfastq-dump, hisat2-mapping, samtools_sam2bam, samtools_sort, stringtie | https://github.com/pitagora-galaxy/cwl/tree/master/workflows/hisat2-stringtie |
| star-cufflinks | download-sra, pfastq-dump, star-mapping, samtools_sam2bam, samtools_sort, cufflinks | https://github.com/pitagora-galaxy/cwl/tree/master/workflows/star-cufflinks |
| star-stringtie | download-sra, pfastq-dump, star-mapping, samtools_sam2bam, samtools_sort, stringtie | https://github.com/pitagora-galaxy/cwl/tree/master/workflows/star-stringtie |
| kallisto | download-sra, pfastq-dump, kallisto-quant | https://github.com/pitagora-galaxy/cwl/tree/master/workflows/kallisto |
| salmon | download-sra, pfastq-dump, salmon-quant | https://github.com/pitagora-galaxy/cwl/tree/master/workflows/salmon |

Table 4 shows that the summary of runtime metrics, processing duration, and the calculated cost of instance usage per run for two workflows, HISAT2-Cufflinks and TopHat2-Cufflinks. The fastest processing time was one of the HISAT2-Cufflinks workflow run on the c5.4xlarge instance, but the execution at the cheapest cost was the HISAT2-Cufflinks workflow on the c5.2xlarge instance. It indicates that workflows on cloud instances can have a trade-off of the processing time and the financial cost. The priority of the research project, the execution speed over the financial cost or vice versa, will be required for the final decision of instance selection optimization. The table also shows the possibility of loss of time or money when one failed to choose a proper instance type. For example, if one used the r5.4xlarge instance to run the HISAT2-cufflinks workflow, it is 7% slower than c5.4xlarge, and about 1.6 times expensive per sample. The impact of the instance type optimization failure will be more serious for the data processing jobs that take days or weeks.

Figure 4 shows the results of processing duration of the HISAT2-StringTie workflow. There are clear differences of processing time between the samples, where the samples of the smaller number of reads have smaller differences between the instance types, but the runs on instance types with more CPU (4xlarge) marked shorter processing time with the samples of the larger number of reads. Each workflow runs used as many CPU cores as available on the environment; thus the difference can be considered as the difference of the number of threads. The read length and the processing duration also have a strong linear relationship. This result will be useful to estimate the resource usage from the size of input data. Supplementary Figure 1 shows the plots of the processing time of the different workflows in which the similar results were shown.

**Figure 4:**
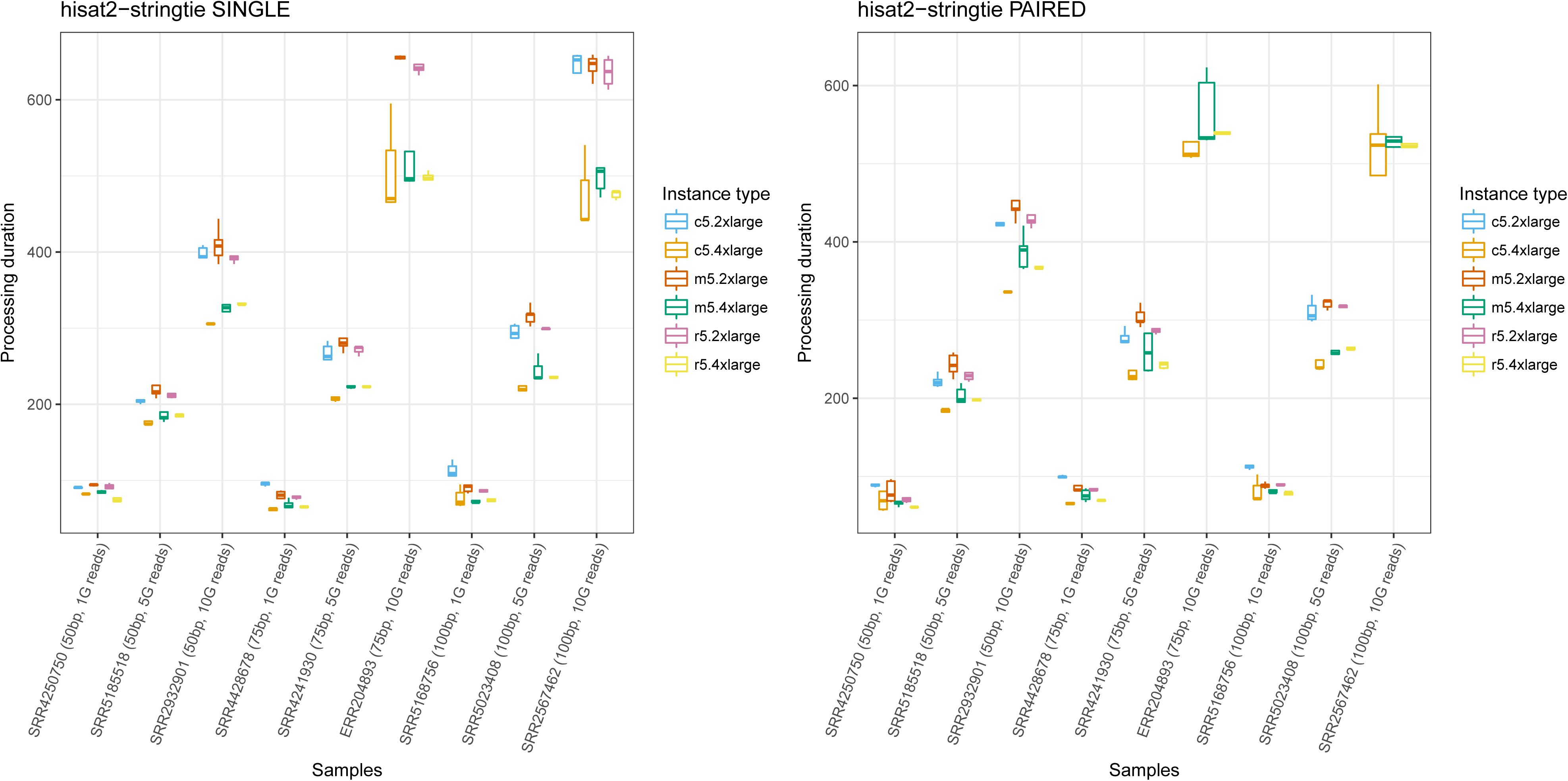
Box plot of per sample processing duration distribution of HISAT2-StringTie workflow. We plotted the values of processing duration of workflow runs excluding data download time. The x-axis shows SRA Run ID of samples used as input data with read length and number of reads. The y-axis shows the workflow processing duration in seconds. Values are separated and colored by the used instance type. Some runs on specific instance types are not in the plots because the failed executions are excluded. Each combination of sample and instance type were iterated five times to show the distribution of metrics. The plot shows that read length and the number of reads are both the factors that effect to the processing duration, and the differences between instance types are relatively small with the smaller number of reads (1G bases), while instances with more CPU cores (*.4xlarge) show shorter processing duration with 10GB reads.

On the other hand, the result of the comparison of the total amount of memory per input data in Supplementary Figure 2 needs a different interpretation. Unlike HISAT2 and TopHat2, Kallisto and Salmon did not show clear differences in memory usage in different sizes of input data. The result indicates that the users need to know the behavior of the tool beforehand since the resource usage depends on the algorithms and the implementations.

The runtime metrics data provided by CWL-metrics also helps to perform a tool comparison. Figure 5 shows that the difference of processing time between the used workflows. Although users need to know the difference of the design concept and the strength of the tools to select the proper one for their research objectives, this result helps to understand the difference of the resource requirement of the workflows for similar purpose. For example, HISAT2 and STAR marked almost the same processing time, but STAR uses far more amount of memory. The plot of the processing time also shows that the obsolete tool TopHat2 is remarkably slower than the other tools.

**Figure 5:**
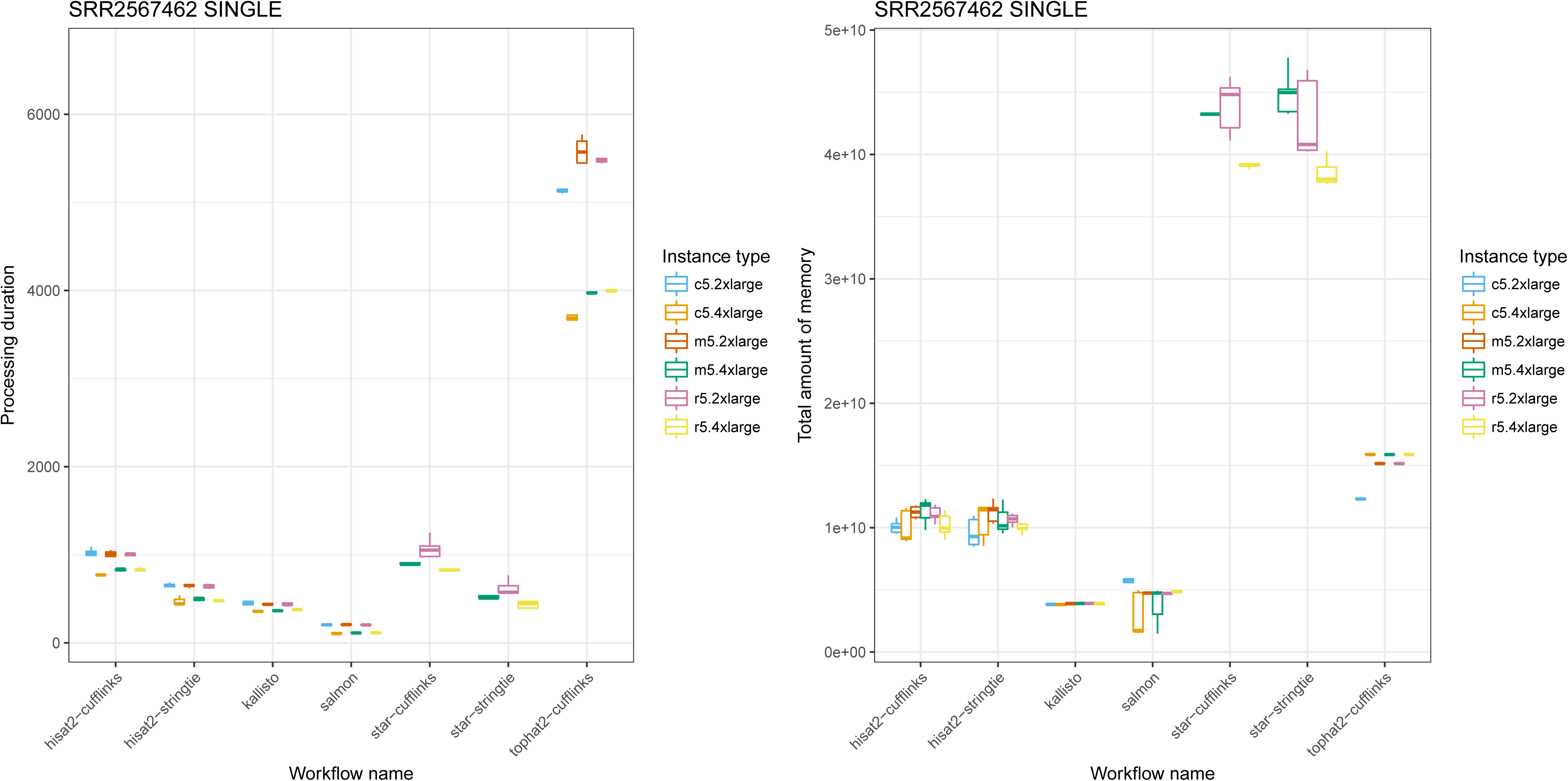
Box plot of processing duration and maximum memory usage of sample SRR2567462 per workflow. The values of processing duration were without data download time. Both plots used values of workflow executions as single end input of SRR2567462. The x-axis shows workflow names, and the y-axis shows the processing duration in seconds and total memory usage in bytes. We iterated each combination of workflow and instance type for five times. The plot of processing duration shows that there is a significant difference in execution time between the TopHat2 workflow and the others. While the difference of processing durations is relatively small, workflows with STAR aligner require four or five times much memory than HISAT2 workflows. These data suggest users know about runtime metrics of workflows before selecting cloud instance type.

## Discussion

CWL-metrics enabled users to choose a proper cloud instance for workflow runs based on the runtime metrics data. The metrics data summarized by workflow inputs, such as the number of threads to use or total file size of input data, provides the most efficient cloud use for a research project. The data will also help the administrator of computational infrastructure to encourage researchers to use the cloud environment in case their local environment has too many running jobs to accept new job submissions. Each user might perform different analyses and visualizations concerning input parameters of their interest. Thus CWL-metrics outputs JSON and TSV data which are easy to parse and used for visualization by any language of users’ favorite, rather having a custom visualization tool other than Kibana.

CWL-metrics is applicable for most cases in bioinformatics data analysis. However, there are cases that the system does not work as effectively as expected. For example, the current implementation of CWL-metrics cannot capture the precise runtime metrics data of a tool that scatter its processes to multiple computation nodes. Also, it cannot estimate the performance of software that uses hardware acceleration systems such as GPU, since the information of those specific architectures is not available via Docker API. Nevertheless, in the example use case using RNA-Seq workflows, we showed CWL-metrics could provide beneficial information to help users to decide on the use of cloud infrastructure.

There are also the other workflow operation frameworks that have functions to capture runtime metrics, such as Galaxy [5], Toil [10], or Nextflow [11]. However, we chose CWL as the workflow description framework and its reference implementation cwltool as the workflow runner for the system because CWL is the project providing a way to share the workflow across the different workflow systems. Once users collected the runtime metrics of workflows by CWL-metrics, they can use the same workflow description with multiple workflow runner implementations. There are fifteen implementations listed as those supporting CWL [12]. Some implementations including Galaxy are still not covering full functions to import and export CWL description to share and run workflows, but the others including Arvados, Toil, and Apache Airflow are already available to users. If one wanted to use a workflow system that does not support CWL yet, the summary of runtime metrics collected through Docker container is still valuable resource across the different frameworks.

CWL project has a subproject, CWL-Prov, to provide the provenance information of workflow executions to improve reproducibility of workflows by tracking intermediate files and logs [13]. The provenance information helps users to track inputs and outputs of workflow runs by using file checksum but does not record the detail of the resource usage. Adding runtime metrics data into the provenance information will cover the information regarding deployment, which helps users to reproduce the runs on a proper computing environment. Thus, the summary of runtime metrics collected by CWL-metrics should be bundled with the provenance information.

There will be more amount of sequencing data that one researcher needs to process by the technologies that produce a large amount of sequencing data such as high-throughput single-cell sequencing. In such a situation, it is essential to have a flexible computing environment that can quickly scale out according to the amount of data. The fast deployment of the data analysis environment to the proper cloud instance supported by Docker, CWL, and CWL-metrics is a way to achieve the computational scale out, which brings a huge benefit for bioinformatics researchers.

### Potential Implications

The Common Workflow Language project aims to provide the workflow description specification for all domains that work with data analysis pipelines.Therefore, CWL-metrics can contribute to other domains through the application of CWL. Sharing CWL workflows with the metrics data captured by CWL-metrics can help users to deploy them on an appropriate environment.

## Methods

### CWL-metrics software components

CWL-metrics runtime metrics capturing system is composed of five software components: Telegraf [14], Fluentd [15], Elasticsearch [16], Kibana [ 17], and a Perl daemon script. Telegraf is an agent to collect runtime metrics of running containers via Docker API using Telegraf Docker plugin. Fluentd works as a log data collector to send metrics data produced by Telegraf to Elasticsearch server. Elasticsearch is a data store to accumulate runtime metrics data and workflow metadata, accepting JSON format data via API endpoint. Kibana is a data browsing dashboard for Elasticsearch to view raw JSON data and to summarize and visualize data. Telegraf, Fluentd, Elasticsearch/Kibana launch as a set of containers during the initialization of CWL-metrics. CWL-metrics runs a Perl script which monitors processes on the host machine to capture cwltool processes. Once the script found a cwltool process, the script runs a function to collect workflow information via debug output of the cwltool process, "docker info" command output, Docker container log via "docker ps" command, and output of system commands to collect environment information. CWL-metrics provides a command *cwl-metrics*, which allows users to start and stop the metrics collection system, and fetch summarized runtime metrics data in a specified format, JSON or tab-separated format. The script to launch the whole system, CWL-metrics installation instruction, and the documentation are available on GitHub [18].

### Packaging RNA-Seq tools and workflows

We used seven different RNA-Seq quantification workflows to capture runtime metrics and analyze performance on cloud infrastructure. Each workflow starts with the tool to download sequence data from Sequence Read Archive (SRA), then convert SRA format file to FASTQ format. Consequently, each pipeline does sequence alignment to reference genome sequence (HISAT2, STAR, and TopHat2) or alignment-like approaches (Kallisto and Salmon) to the set of reference transcript sequence, then perform transcript quantification. Most of the tool containers used in the workflows are from the Biocontainers [19] registry. We containerized the tools those are not available on the registry and uploaded them to the container registry service Quay [20]. We described tool definitions such as input and output of tool execution and the workflow procedures in CWL tool files, which are available on GitHub [21]. Each workflow has two options for sequence read layout single-end and paired-end; thus we used fourteen workflows in total. The Supplementary Table 1 shows the tool versions, the online location of the CWL tool files, and the original tool website locations.

### Select RNA-Seq workflow input sequence data from the public data repository

To analyze the effect of sequence data quality to workflow runtime performance, we chose nine samples of different read length and number of reads from the public raw sequencing data repository, SRA (Table 2). We used the Quanto database [22] to select the data by filtering length and number of sequence reads, with the condition of read length, 50, 75, or 100 and the approximate number of sequence, 1,000,000, 5,000,000, or 10,000,000. We filtered the data with the query "organism == Homo sapiens", "study type == RNA-Seq", "read layout == PAIRED", and "instrument model == Illumina HiSeq", then manually picked suitable data. Both single-end and paired-end workflows used the same dataset while single-end workflows treated paired-end read files reads as two single-end read files. The version of the reference genome is GRCh38. We downloaded the reference genome file from the UCSC genome browser [23], and the transcriptome was from Gencode [24].

**Table 2:** The read characteristics of processed RNA-Seq data. We chose nine different RNA-Seq data from the SRA, a public high-throughput sequencing data. Each data are different in their read length and a total number of reads for performance comparison. All data are from human sample sequenced by the Illumina HiSeq platform.

| SRA Run ID | Read length | Number of reads per strand | BioSample ID | Sample description | Sequencing instrument |
| --- | --- | --- | --- | --- | --- |
| SRR4250750 | 50 | 1,000,425.00 | SAMN05779985 | cultured embryonic stem cells | Illumina HiSeq 2500 |
| SRR5185518 | 50 | 5,008,398.00 | SAMN06239034 | cultured embryonic stem cells | Illumina HiSeq 2500 |
| SRR2932901 | 50 | 10,017,495.00 | SAMN04211783 | fetal lung fibroblasts | Illumina HiSeq 2500 |
| SRR4428678 | 75 | 1,043,870.00 | SAMN05913930 | embryonic stem cell derived macrophage | Illumina HiSeq 4000 |
| SRR4241930 | 75 | 5,004,985.00 | SAMN05770731 | PGC-like cells (PGCLCs) | Illumina HiSeq 2000 |
| ERR204893 | 75 | 10,234,883.00 | SAMEA1573291 | lymphoblastoid cell line | Illumina HiSeq 2000 |
| SRR5168756 | 100 | 1,006,868.00 | SAMN06218220 | subcutaneous metastasis | Illumina HiSeq 2500 |
| SRR5023408 | 100 | 5,004,554.00 | SAMN06017954 | primary breast cancer | Illumina HiSeq 2500 |
| SRR2567462 | 100 | 10,007,044.00 | SAMN04147557 | prostate cancer cells LNCaP | Illumina HiSeq 2500 |

### Run workflows on AWS EC2

To evaluate the performance on running different RNA-Seq workflows, we selected instance types of two different sizes 2xlarge and 4xlarge from three categories, general purpose, compute optimized, and memory optimized to run all workflows for all samples (Table 3). Each combination of instance type, workflow, and sample data was executed for five times while CWL-metrics is running on the same machine to capture the runtime metrics information. All workflow runs used Elastic Block Storage of General Purpose SSD volumes as file storage. We downloaded all the reference data used for workflows in advance. The scripts to get reference data and run workflows are available online [21].

**Table 3:** The machine specs of AWS EC2 instance types used in the metrics collection. To compare the performance of workflow runs on different computing platforms, we selected three categories from AWS EC2 categories, general purpose, compute optimized, and memory optimized. We further selected two different instance types from those three categories according to the number of virtual CPUs, 2xlarge and 4xlarge, with 8 and 16 CPU cores, respectively. Instance usage prices are as of 14 August 2018 for on-demand use in the US East (N. Virginia) region. Prices are not including charges for storage, network usage, and other AWS features.

| Instance type | Category | vCPU | ECU | Memory (GiB) | Linux/UNIX Usage (per Hour) |
| --- | --- | --- | --- | --- | --- |
| m5.2xlarge | General Purpose | 8 | 31 | 32 | \$0.384 |
| m5.4xlarge | General Purpose | 16 | 60 | 64 | \$0.768 |
| c5.2xlarge | Compute Optimized | 8 | 34 | 16 | \$0.34 |
| c5.4xlarge | Compute Optimized | 16 | 68 | 32 | \$0.68 |
| r5.2xlarge | Memory Optimized | 8 | 31 | 64 | \$0.504 |
| r5.4xlarge | Memory Optimized | 16 | 60 | 128 | \$1.008 |

**Table 4:**
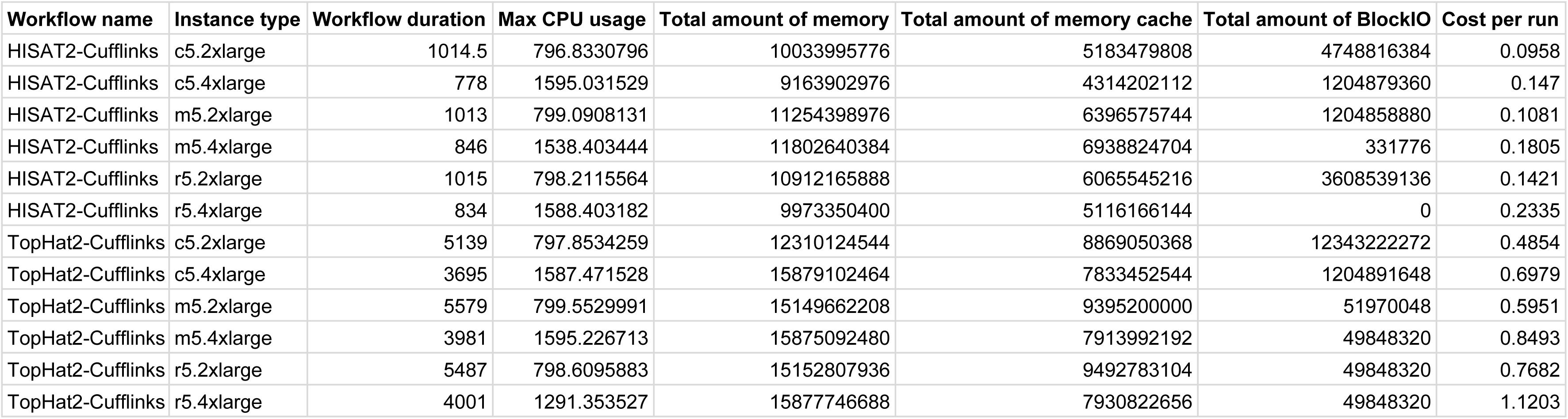
The runtime metrics comparison of TopHat2 and HISAT2. We summarized the runtime metrics values to compare two different workflows HISAT2-cufflinks and TopHat2-cufflinks. All runs are of input data SRR2567462. The read length was 100bp, the number of reads was 10,007,044.00, and the read layout was single-end. The shown values are workflow duration in seconds, the maximum CPU usage in percentage, the total amount of memory in bytes, the total amount of cache in bytes, the total amount of block IO in bytes, and the cost per run in USD. We calculated the median values for metrics values from the data of five times workflow iteration. Values can be zero for short-time steps since CWL-metrics collects these metrics with sixty seconds interval.

### Collect runtime metrics and summarize

After the workflow executions, we collected summarized metrics data from Elasticsearch by *cwl-metrics fetch* command. Exported JSON format data were parsed by a ruby script to create data summarized per workflow runs, loaded on Jupyter notebook [25] for further analysis. We calculated statistics of metrics by R language functions [26], and we created the box plots by the ggplot2 package [27]. The notebook file is available on GitHub [28].

## Figure legends

**Supplementary Figure 1: Box plot of processing duration for all workflows** The x-axis shows SRA Run ID of input data with the read length and the number of reads. The y-axis shows the processing duration in seconds excluding data downloading time. In most of the used workflows, the read length and the number of reads of input data affect the processing time. Workflows with STAR aligner requires a large amount of memory; thus the executions on instance types with a smaller amount of memory have failed.

**Supplementary Figure 2: Box plot of max memory usage for all workflows** The x-axis shows SRA Run ID of input data with the read length and the number of reads. The y-axis shows the maximum amount of memory used during the process in bytes. The distributions of values are large especially on runs which finishes in a short time because sixty seconds interval of metrics capturing could not get the right values.

**Supplementary Table 1: The versions and containers of tools used in the RNA-Seq workflows**

We used eleven tools in total to construct seven RNA-Seq quantification workflows. The two tools we developed, download-sra and pfastq-dump, are packaged in containers by ourselves. The container of Salmon was available on its developer’s build. We found the rest of tools in Biocontainers registry. We wrapped all the tools as CWL CommandLineTool class files and available on GitHub.

## Availability of source code and requirements

For CWL-metrics, the runtime metrics capturing system:

Project name: cwl-metrics

Project home page: https://inutano.github.io/cwl-metrics/

Operating system(s): Platform independent

Programming language: Perl v5.18.2 or higher

Other requirements: Docker 18.06.0-ce or higher and Docker Compose 1.22.0 or higher, cwltool 1.0.20180820141117 or higher

License: MIT

Any restrictions to use by non-academics: NA

For the scripts and the notebook for visualization on this manuscript:

Project name: cwl-metrics-manuscript

Project home page: https://github.com/inutano/cwl-metrics-manuscript

Operating system(s): Platform independent

Programming language: Ruby 2.5.1 or higher

Other requirements: Docker 18.06.0-ce or higher

License: MIT

Any restrictions to use by non-academics: NA

## Availability of supporting data and materials

The data set used for the visualizations of this article is available in figshare [29]. The full summary data and visualization on Jupyter notebook is available on GitHub [30] and nbviewer [31].

## Declarations

### List of abbreviations

CWL: Common Workflow Language
AWS: Amazon Web Service,
TSV: tab separated values
EC2: Elastic Compute Cloud
SRA: Sequence Read Archive

### Authors’ contributions

Conceptualization, Methodology, Software, Investigation: TO TT. Visualization, Writing original draft: TO. Supervision: OO.

## Acknowledgements

The authors are grateful to Prof. Kento Aida and the Inter-Cloud CREST team for constructive comments and discussions. The authors also thank the open source communities: Pitagora Galaxy, Galaxy Project, Common Workflow Language, Bioinformatics Open Source Conference, and the BioHackathon for many comments and suggestions. We performed implementation and testing of the system on the NIG supercomputer at ROIS National Institute of Genetics.

## References

1. Chang J. Core services: Reward bioinformaticians. Nature 2015;520:151–2.

2. Prins P, de Ligt J, Tarasov A et al. Toward effective software solutions for big biology. Nature Biotechnology 2015;33:686–7.

3. Merkel D. Docker: lightweight linux containers for consistent development and deployment. Linux Journal. 2014 Mar 1;2014(239):2.

4. Di Tommaso P, Palumbo E, Chatzou M et al. The impact of Docker containers on the performance of genomic pipelines. PeerJ 2015;3:e1273.

5. Afgan E, Baker D, Batut B et al. The Galaxy platform for accessible, reproducible and collaborative biomedical analyses: 2018 update. Nucleic Acids Research 2018;46:W537–44.

6. Amstutz P, Crusoe MR, Nebojša Tijanić et al. Common Workflow Language, v1.0. 2016, DOI: 10.6084/m9.figshare.3115156.v2.

7. Stein LD. The case for cloud computing in genome informatics. Genome Biology 2010;11:207.

8. Amazon EC2 Instance Types https://aws.amazon.com/ec2/instance-types/ Accessed 30 Oct 2018.

9. common-workflow-language/cwltool https://github.com/common-workflow-language/cwltool Accessed 30 Oct 2018.

10. Toil: A scalable, efficient, cross-platform pipeline management system written entirely in Python and designed around the principles of functional programming. http://toil.ucsc-cgl.org/ Accessed 30 Oct 2018.

11. Di Tommaso P, Chatzou M, Floden EW et al. Nextflow enables reproducible computational workflows. Nature Biotechnology 2017;35:316–9.

12. Common Workflow Language https://www.commonwl.org/ Accessed on 30 Oct 2018.

13. Khan FZ, Soiland-Reyes S, Sinnott RO et al. CWLProv: Interoperable Retrospective Provenance Capture And Computational Analysis Sharing. 2018, DOI: 10.5281/zenodo.1473157.

14. Telegraf https://www.influxdata.com/time-series-platform/telegraf/ Accessed 30 Oct 2018.

15. Fluentd https://www.fluentd.org/ Accessed 30 Oct 2018.

16. Elasticsearch https://www.elastic.co/products/elasticsearch Accessed 30 Oct 2018.

17. Kibana https://www.elastic.co/products/kibana Accessed 30 Oct 2018.

18. CWL-metrics https://inutano.github.io/cwl-metrics/ Accessed 30 Oct 2018.

19. da Veiga Leprevost F, Grüning BA, Alves Aflitos S et al. BioContainers: an open-source and community-driven framework for software standardization. Valencia A (ed.). Bioinformatics 2017;33:2580–2.

20. QUAY - inutano https://quay.io/user/inutano Accessed 30 Oct 2018.

21. pitagora-galaxy/cwl https://github.com/pitagora-galaxy/cwl Accessed 30 Oct 2018.

22. Ohta T, Nakazato T, Bono H. Calculating the quality of public high-throughput sequencing data to obtain a suitable subset for reanalysis from the Sequence Read Archive. GigaScience 2017;6, DOI: 10.1093/gigascience/gix029.

23. Casper J, Zweig AS, Villarreal C, Tyner C, Speir ML, Rosenbloom KR, Raney BJ, Lee CM, Lee BT, Karolchik D, Hinrichs AS. The UCSC genome browser database: 2018 update. Nucleic acids research. 2017;46(D1):D762-9.

24. Harrow J, Frankish A, Gonzalez JM et al. GENCODE: The reference human genome annotation for The ENCODE Project. Genome Research 2012;22:1760–74.

25. Project Jupyter http://jupyter.org Accessed 30 Oct 2018.

26. R Core Team. R: A language and environment for statistical computing. R Foundation for Statistical Computing, Vienna, Austria. 2015. https://www.R-project.org/. Accessed 30 Oct 2018.

27. H. Wickham. ggplot2: Elegant Graphics for Data Analysis. Springer-Verlag New York; 2009.

28. inutano/cwl-metrics https://github.com/inutano/cwl-metrics Accessed 30 Oct 2018.

29. Ohta, T. Runtime metrics data of 7 different RNA-Seq quantification workflows. figshare. 2018-10-18 https://doi.org/10.6084/m9.figshare.7222775.v1

30. inutano/cwl-metrics-manuscript https://github.com/inutano/cwl-metrics-manuscript Accessed 30 Oct 2018.

31. CWL-metrics: workflow runtime metrics analysis https://nbviewer.jupyter.org/github/inutano/cwl-metrics-manuscript/blob/master/notebook/CWL-metrics%20runtime%20metrics%20analysis.ipynb Accessed 30 Oct 2018.

